# Profiling blood-based neural biomarkers and cytokines in experimental autoimmune encephalomyelitis model of multiple sclerosis using single-molecule array technology

**DOI:** 10.1101/2023.12.25.573285

**Authors:** Insha Zahoor, Sajad Mir, Shailendra Giri

**Affiliations:** Department of Neurology, Henry Ford Health, Detroit, MI 48202, USA

**Keywords:** Biomarker, Cytokines, Glial Fibrillary Acidic Protein (GFAP), Neurofilament Light Chain (NFL), Single Molecule Array (SIMOA), Experimental Autoimmune Encephalomyelitis (EAE), Multiple Sclerosis (MS)

## Abstract

Experimental autoimmune encephalomyelitis (EAE) is a preclinical animal model widely used to study multiple sclerosis (MS). Blood-based cytokines and neural biomarkers are predictors of neurodegeneration, disease activity, and disability in patients with MS. However, understudied confounding factors cause variation in reports on EAE across animal strains/studies, limiting the utility of these biomarkers for predicting disease activity. In this study, we investigated blood-based analyte profiles, including neural markers (NFL and GFAP) and cytokines (IL-6, IL-17, IL- 12p70, IL-10, and TNF-α), in two clinically distinct EAE models: relapsing-remitting (RR)-EAE and chronic-EAE. Ultrasensitive single-molecule array technology (SIMOA, Quanterix) was used to profile the analytes in blood plasma of mice at the acute, chronic, and progressive phases of disease. In both models, NFL was substantially increased during post-disease onset at the peak, chronic, and progressive phases, with pronounced increase in chronic-EAE. Leakage of GFAP into peripheral blood was also greater after disease onset in both EAE models, especially in the acute phase of chronic-EAE. Among all cytokines, only IL-10 had consistently lower levels in both EAE models throughout the course of disease. This study suggests NFL, GFAP, and IL-10 as potential translational predictors of disease activity in EAE, making them potential candidates for surrogate markers for preclinical testing of therapeutic interventions in animal models of MS.

## 1. Introduction

Multiple sclerosis (MS) is an inflammatory disease of the central nervous system (CNS) characterized by immune-mediated damage to the insulating myelin sheath, resulting in progressive demyelination and neurodegeneration. Experimental autoimmune encephalomyelitis (EAE) is a well-established preclinical animal model of MS (Gold et al., 2006; Steinman and Zamvil, 2005; 2006; Farooqi et al., 2010; Bjelobaba et al., 2018). Despite certain inherent limitations, the development of potential treatment options for MS has improved (Steinman, 2009; Steinman et al., 2023). EAE represents a complex heterogeneous model that relies on the disease induction method used, which adds variability to disease pathogenesis, making it an imperfect but approximate model for studying MS. It is worth mentioning that its use depends entirely on the scientific question addressed and the feasibility of translation. There are two widely used EAE models that approximate the relapsing-remitting (RR) and chronic disease phenotypes of MS: the SJL-based biphasic RR-EAE model, and the monophasic B6-chronic model (Baker and Amor, 2015; 2014; Amor and Baker, 2012; Miller et al., 2007). The disease is induced in EAE through an immune-mediated response to CNS components by active immunization with self-antigens to myelin, making it an inflammatory model (Constantinescu et al., 2011; McCarthy et al., 2012). The CNS antigens targeted are myelin basic protein (MBP), myelin oligodendrocyte glycoprotein (MOG), or proteolipid protein (PLP). EAE, as an animal model, is considered the backbone for preclinical predictions and testing potential drugs for MS by monitoring effects on histopathology, motor deficits, and inflammatory responses in EAE (Constantinescu et al., 2011). Therefore, it continues to be the most appropriate and relevant animal model to study MS 90 years since its inception (Steinman et al., 2023).

EAE is characterized by impaired inflammation resolution accompanied by dysregulation of cytokine profiles, immune cell infiltration into the CNS, disruption of the blood–brain barrier (BBB), neuroinflammation, and neuroaxonal pathologies, including demyelinating lesions and gliosis (Kipp et al., 2017). Although the precise mechanism of EAE pathogenesis is still unclear, the pathogenic cascade mainly involves the production of cytokines by immune cells throughout the disease course, ultimately compromising BBB integrity and tissue damage (Constantinescu et al., 2011). Measuring blood-based markers is developing importance in assessing the disease activity of MS and its animal model EAE (Birmpili et al., 2022). Effective biomarkers for MS diagnosis, disease progression, and treatment responses, particularly those measurable in the blood, are sorely lacking. While significant effort has been made to identify biomarkers from cerebrospinal fluid (CSF) to diagnose MS, this endeavor has been challenging with limited success (Momtazmanesh et al., 2021). The analysis of easily drawn biofluids, including blood for MS biomarkers, has been minimally investigated but holds significant promise. Neurofilament light (NF-L), glial fibrillary acidic protein (GFAP), and cytokines (IL-17, IL-6, IL-10, TNF-α, IFN-γ, and IL-12) are emerging biomarkers for determining inflammatory disease activity, disease progression, treatment response, and prognosis in MS patients (Jahan-Abad et al., 2020; Kuenz et al., 2007; Göbel et al., 2018; Imitola et al., 2005; Palle et al., 2017). The leakage of NFL and GFAP into plasma/serum is particularly useful for monitoring neuroaxonal damage in MS, and yet there are limited studies on this topic in preclinical animal models of EAE (Jahan-Abad et al., 2020; Aharoni et al., 2021; Gnanapavan et al., 2012; Brummer et al., 2023).

Given the urgent need for blood-derived biomarkers for MS, the inception of highly sensitive immunoassays serves as the foundation. Conventional methods of measuring blood biomarkers have certain limitations due to low sensitivity and inconsistency across reports showing profiles of neural markers and cytokines in preclinical models (Kuhle et al., 2016). Comparisons across profiling studies of EAE often lack reproducibility due to variability across mice and underlying confounding factors such as the extent of inflammation or demyelination and small sample sizes (Aharoni et al., 2021). The biological matrix used for profiling blood-based analytes also contributes to variability.

With the increase in neuroprotective or reparative strategies, potential therapeutics for EAE must be evaluated through quantitative neuropathology. Herein, we applied SIMOA (SIngle MOlecule Array), as this is an emerging ultrasensitive technology based on a bead-conjugate immunocomplex with the potential to detect analytes present at ultralow levels (femtomolar range) and below detection limits for other conventional assays (Cohen et al., 2020; Lasseter et al., 2020; Revendova et al., 2022; Pafiti et al., 2023). The average number of enzymes per bead (AEB) was considered the unit of measurement. Several different biomarkers can be detected in a single experiment, in a variety of sample types, with singleplex and multiplex assays. According to Quanterix, this digital platform can precisely measure analyte levels in human blood samples with 1,000 times greater sensitivity than conventional measurements (https://www.quanterix.com/simoa-technology/). This means that subtle changes can be captured earlier and less invasively compared to other approaches. In this study, we aimed to further investigate the outcome of EAE manifestations in the form of disease severity parameters and detectability of plasma NFL, GFAP, and cytokines over the course of the disease. Overall, we propose the application of SIMOA as a valuable and useful approach for indirect assessments of disease pathology and CNS damage in EAE.

## 2. Materials and Methods

### EAE model induction

A group of 5-10 female SJL/J and C57BL/6 (ten- to twelve-week-old) mice were purchased from the Jackson Laboratory (Bar Harbor, ME). Animals were housed in a pathogen-free animal facility at Henry Ford Health, Detroit, MI, according to animal protocols approved by IACUC. All mice were housed with standard food and water ad libitum at a room temperature of 22 ± 2°C under a 12:12 h light-dark cycle. For the RR-EAE model, SJL mice were immunized on day 0 via subcutaneous injections in the flank region with the antigen PLP_139-151_ peptide (100 μg/mouse) emulsified in 200 uL Complete Freund’s Adjuvant (CFA) (Sigma Chemicals, St. Louis, MO, USA) supplemented with 4 mg/ml heat-killed *Mycobacterium tuberculosis* H37Ra (400 μg; Becton, Dickinson and Company, Sparks, MD, USA) as described previously (Mangalam et al., 2013; Mangalam et al., 2016; Poisson et al., 2015; Zahoor et al., 2022; Zahoor et al., 2025). For the chronic model, EAE was induced in B6 mice on day 0 with 200 uL CFA containing the antigen MOG_35-55_ (200 µg/mouse) prepared similarly, as described previously (Mangalam et al., 2013; Mangalam et al., 2016; Poisson et al., 2015). Pertussis toxin (List Biological Laboratories, CA, USA) was administered via intraperitoneal (i.p.) injection (200 ng/200 μL) on days 0 and 2. In both models, one set of mice was injected with CFA without antigen/peptide to serve as a control.

### Clinical assessment of EAE models

For clinical evaluation of models based on neurological deficits, the animals were monitored daily and clinical score was recorded, until the duration of the study, in a blinded fashion by observing disease signs, including paralysis, according to the conventional grading system on a scale of 0- 4, as described previously (Mangalam et al., 2013; Mangalam et al., 2016; Poisson et al., 2015; Zahoor et al., 2022; Zahoor et al., 2025). The onset of disease in EAE was considered on the day animals presented the first signs of disease. The disease peak was considered when the EAE score reached its maximum and did not increase from the previous day (16–19 days postimmunization). The chronic phase of the disease was considered after 20 days postimmunization if the EAE score decreased, or was maintained, after the disease peak. Weight loss was monitored after EAE induction. Changes in weight during the disease course were evaluated for each mouse. Readings were taken every week. Animals were provided appropriate supportive care and maintenance while showing disease signs. Mice with severe condition called for immediate premature euthanasia to meet humane endpoint criteria.

### Experimental groups and blood sampling

Mice were euthanized at different time points during the disease course. The time points selected for EAE profiling included day 17 (peak), day 30 (chronic), and day 45 (progressive) for RR-EAE (SJL) and day 15 (before peak), day 20 (peak) and day 45 (chronic) for chronic-EAE (B6). Mice were anesthetized with CO_2_ and transcardially perfused with 1X chilled PBS. Control CFA-treated mice were also included at the indicated time points. Blood samples from mice were collected in ethylenediaminetetraacetic acid (EDTA)-containing tubes for plasma analysis. Plasma was separated from blood by centrifugation at 1500 × g for 10 min. The clear yellow liquid supernatant was collected from the top and stored in respective tubes at -80°C until further processing (zero freeze/thaw cycles). The secondary method of euthanasia for mice was either cervical dislocation or needle-induced pneumothorax. All animal experiments in this study were performed per the policies and guidelines of the IACUC committee at Henry Ford Health under the animal welfare assurance number D16-00090.

### Single Molecule Array (SIMOA) Assay

To characterize profiles of neural markers and cytokines in EAE, we measured NFL and GFAP in plasma samples from CFA and EAE mice in both models (RR and chronic) with a commercially available SIMOA™ Neuro 2-Plex B Advantage Kit (product number: 103520) (Quanterix, MA) on an SR-X analyzer, according to the manufacturer’s instructions. Additionally, cytokine profiling was performed using a mouse Cytokine 5-Plex Kit for IL-6, TNF-A, IL-17, IL-12, and IL-10 (Cat # 107-178-1-AB; Product Number: 85-0441) (Quanterix, MA) on an SP-X analyzer, according to the manufacturer’s instructions. Calibrators supplied with the kits were used as standards. The number of mice used for each assay at a given time point was 4-9. Samples were only loaded after centrifugation at 10,000 rpm for 10 min to prevent lipids in the sample from interfering with the assay. For every 96-well plate-based assay, one batch of reagents was used to minimize the lot-specific variations, as emphasized in the protocol instructions by Quanterix.

### Data analysis

Statistical significance was computed by Mann-Whitney test and all values are presented as the median or mean ± SEM wherever applicable. The figure legends (and wherever applicable) mention statistical significance and power in terms of “n” and *p* values. Comparisons between different time points across disease courses in both models were performed using one-way ANOVA Kruskal-Wallis test. All statistical analyses were performed using GraphPad Prism Software, version 9.2.0 (San Diego, California, USA; www.graphpad.com). An asterisk indicates statistical significance * for *p* ≤ 0.05, ** for *p* ≤ 0.01, *** for *p* ≤ 0.001, and **** for *p* ≤ 0.0001 with n > 4 for all variables.

## 3. Results

### EAE increases NFL and GFAP leakage in the peripheral circulation

The mice injected with respective antigens (PLP_139-151_ and MOG_35-55_) for EAE induction in both RR (SJL) and chronic (B6) models showed classical disease course with established symptoms which was evident from clinical scores (**Fig. 1**). As expected, CFA control mice did not show any clinical symptoms of disease. We have previously shown that histological analyses of spinal cords show an extensive demyelination and antigen specific response in EAE, whereas control mice do not exhibit signs of demyelination and antigen specific response (Poisson et al. 2015). In the present study, neuroaxonal damage in the CNS was assessed by measuring leakage of NFL and GFAP into peripheral circulation over the disease course of EAE. As shown in **Fig. 2** and **Fig. 3**, compared to those in the control CFA group, plasma levels of NFL and GFAP were significantly higher at all time points in both EAE models. The mean NFL level for CFA mice in the RR-EAE model was 31.68 ± 1.93 pg/mL compared to EAE mice, with an average value of 297.45 ± 15.77 pg/mL across all time points when combined (**Fig. 2**). During the peak phase of the disease on day 17, compared to CFA mice, the mean NFL level was 326.42 pg/mL (*p* < 0.05; median 173 pg/ml, clinical score ∼3-4) (**Fig. 2A**) in the induced RR-EAE model, and was greater than other time points (day 30, 272.15 pg/ml, *p* < 0.01, median 255.6 pg/mL; day 45, 298.8 pg/mL, *p* < 0.01, median 238.4 pg/mL) (**Fig. 2B; Fig. 2C**), reflecting a substantial increase compared to control mice, clearly highlighting the pathological manifestations of EAE (**Fig. 2, Fig. 6A**). Overall, NFL levels ranged between 60.43 and 899.25 pg/mL in the RR-EAE model across all time points compared to CFA, which was between 18.35 and 57.75pg/mL, suggesting elevated leakage of NFL in EAE (**Table 1**). However, there was no significant difference across disease stages in RR-EAE. Furthermore, the mean NFL level for the entire disease course reached 598.11 ± 133.83 pg/mL in the chronic-EAE model compared to 17.77 ± 4.08 pg/mL in CFA mice, with time-specific mean values of 533.5 pg/mL in the induced EAE mice on day 15 (clinical score ∼3-4) (*p* < 0.05; median 484.3 pg/mL), 405.48 pg/ml on day 20 (*p* < 0.05; median 289.8 pg/mL), and day 45 (mean value 855.37 pg/mL; *p* < 0.0001; median 517.4 pg/mL) (**Fig. 3; Fig. 5B**). Overall, NFL values in induced chronic-EAE models ranged between 92.47 and 1960.59 pg/mL, whereas NFL values in CFA mice ranged between 9.53 and 34.77 pg/mL (**Table 1**).

**Table 1.** Comparative detection range of analytes in plasma of SJL and B6 models using SIMOA.

| Analyte | Conc range (SJL) (pg/mL) |  | Status in EAE | Conc range (B6) (pg/mL) |  | Status in EAE |
| --- | --- | --- | --- | --- | --- | --- |
|  | CFA | RR-EAE |  | CFA | Chronic-EAE |  |
| **NFL** | 18.35 to 57.75 | 60.43 to 899.25 | Increase | 9.53 to 34.77 | 92.47 to 1960.59 | Increase |
| **GFAP** | 7.22 to 168.70 | 45.65 to 654.52 | Increase | 0.00 to 8.17 | 7.62 to 52.40 | Increase |
| **IL-10** | 12.38 to 95.88 | 0.46 to 33.87 | Decrease | 5.07 to 114.40 | 1.92 to 14.80 | Decrease |
| **IL-6** | Level below low calibrator | Level below low calibrator | Not conclusive | Level below low calibrator | Level below low calibrator | Not conclusive |
| **IL-12p70** | Level below low calibrator | Level below low calibrator | Not conclusive | Level below low calibrator | Level below low calibrator | Not conclusive |
| **TNF-α** | Level below low calibrator | Level below low calibrator | Not conclusive | Level below low calibrator | Level below low calibrator | Not conclusive |
| **IL-17** | Level below low calibrator | Level below low calibrator | Not conclusive | Level below low calibrator | Level below low calibrator | Not conclusive |

**Fig 1.**
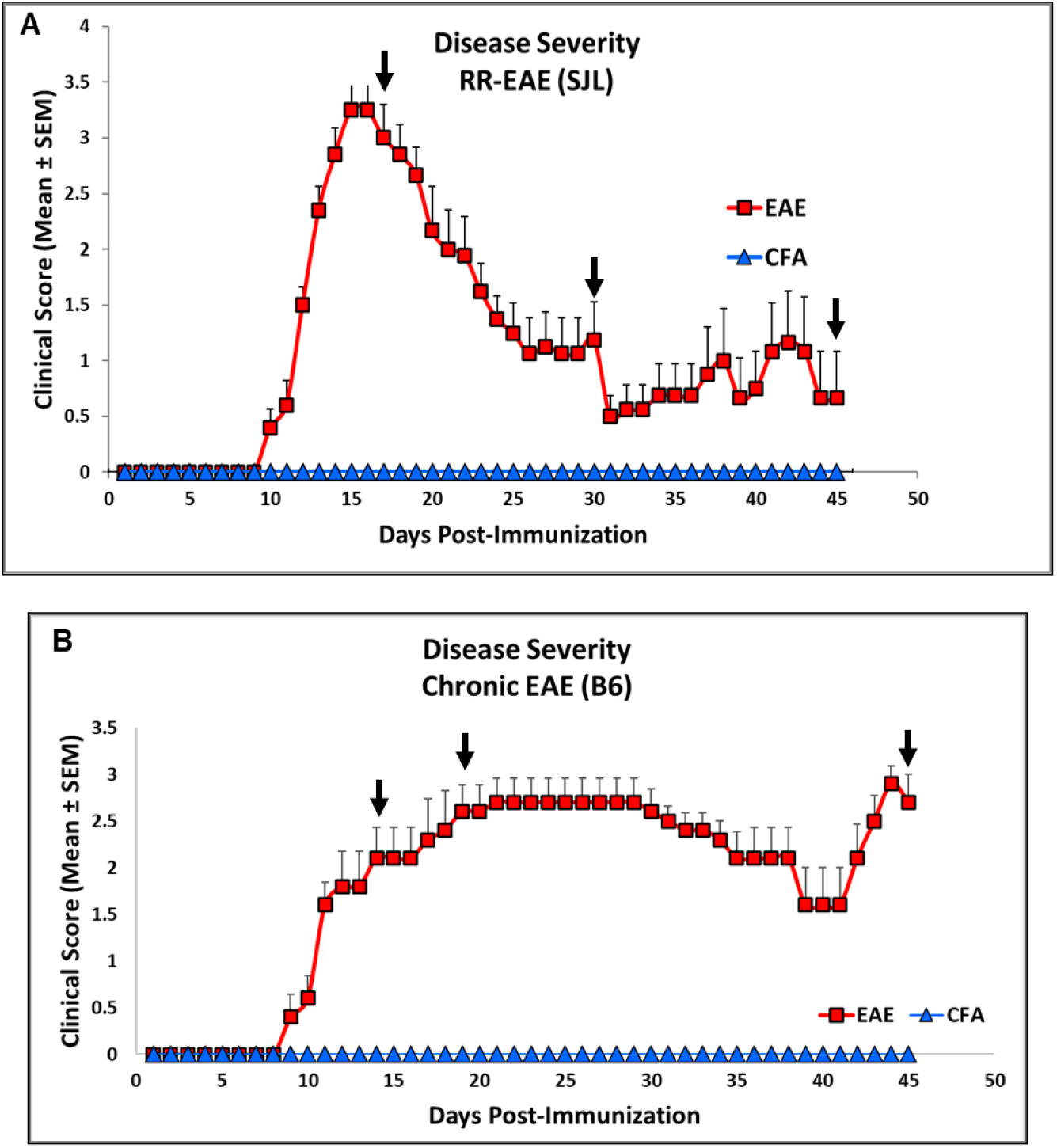
Representative clinical score as indicator of disease severity changes in RR (SJL) and Chronic (B6) model of EAE. (A) Clinical score in SJL and (B) B6 mice consisting of experimental groups CFA and EAE during disease course. Scores are shown as Mean ± SEM.

**Fig 2.**
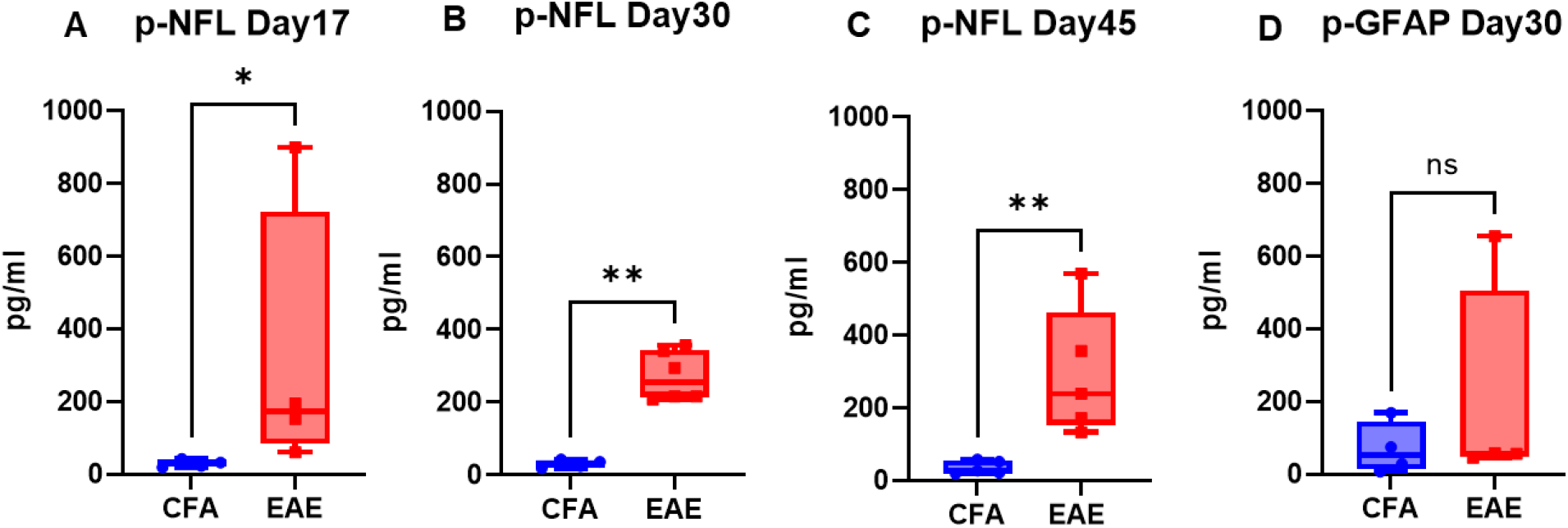
Profile of neural markers (NFL and GFAP) in plasma of RR-EAE. Plasma NFL (A-C) and GFAP (D) levels at respective time points in CFA vs EAE. Values are shown in pg/mL. *p<0.05, **p<0.01 (as determined by Mann-Whitney t-test).

**Fig 3.**
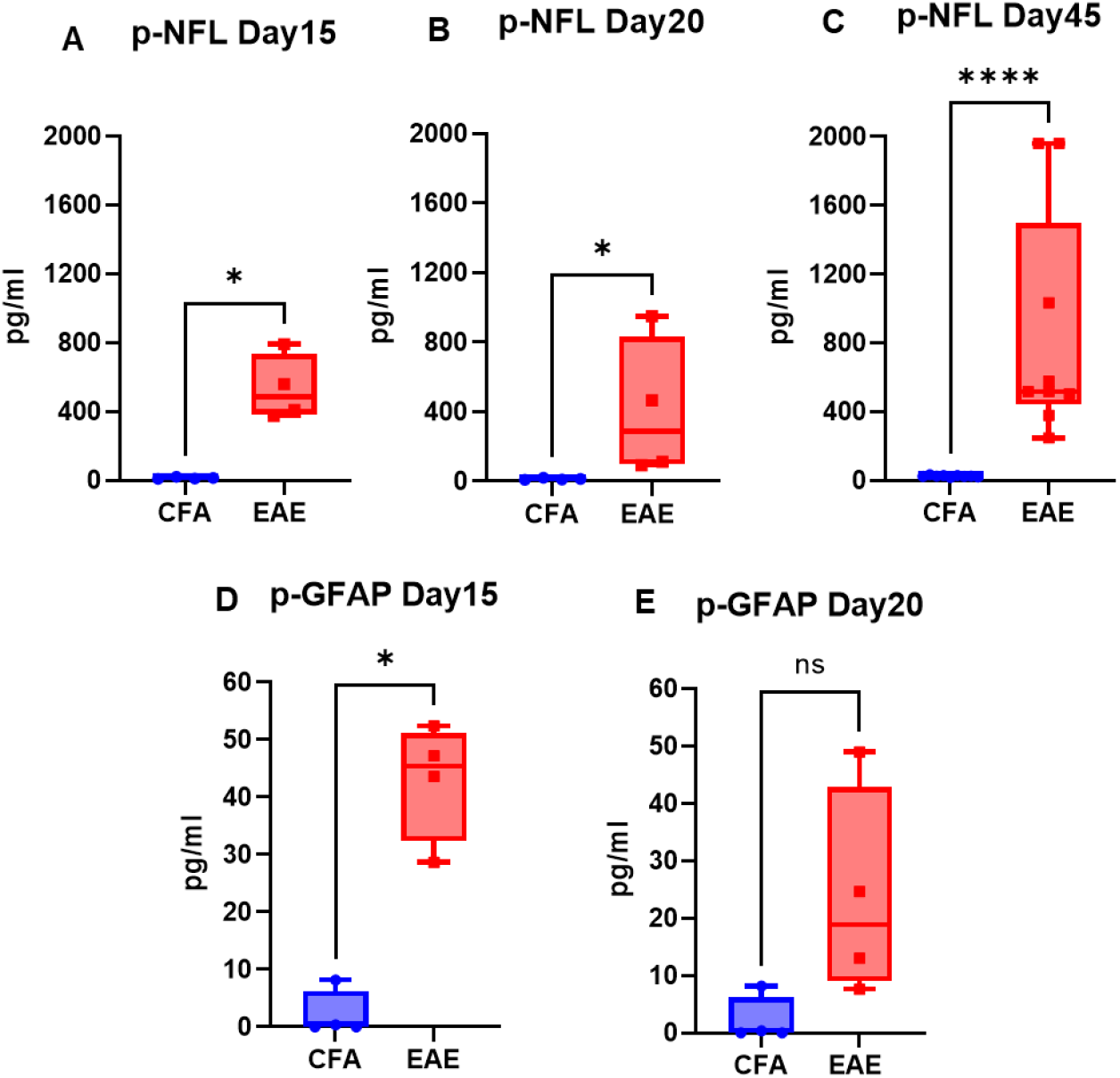
Profile of neural markers (NFL and GFAP) in plasma of Chronic-EAE. Plasma NFL (A-C) and GFAP (D-E) levels at respective time points in CFA vs EAE. Values are shown in pg/mL. *p<0.05, ****p<0.0001 (as determined by Mann-Whitney test).

While the mean plasma level of GFAP in the RR-EAE group was 203.56 ± 173.60 pg/mL (median 57.04 pg/ml), it was 69.88 ± 41.37 pg/ml (median 51.82 pg/ml) in the CFA mice of this model, suggesting that leakage in the EAE induced group was still significant (**Fig. 2D**). The range of GFAP levels was 45.65 to 654.52 pg/mL in induced RR-EAE compared to 7.22 to 168.70 pg/mL in CFA mice (**Table 1**). For the chronic-EAE model, when compared across all time points, the average plasma level of GFAP was 33.27 ± 7.92 pg/mL compared to 2.14 ± 0 pg/mL for CFA. Leakage reached a maximum on day 15 (*p* < 0.05; median 45.43 pg/ml), with an average value near 42.98 pg/mL compared to day 20 with 23.56 pg/mL (median 18.85 pg/ml) in the EAE group (**Fig. 3D, E**). The detection range for GFAP in the chronic-EAE model was 7.62 to 52.40 pg/ml in the induced EAE group compared to 0.00 to 8.17 pg/ml in the CFA group (**Table 1**). Taken together, NFL and GFAP show substantial leakage from the CNS into the periphery in both models of EAE post-disease immunization.

### EAE decreases the level of the anti-inflammatory cytokine IL-10 in the blood

IL-10 plays a crucial role in the pathophysiology of MS and EAE animal models because it is a key cytokine involved in regulating the immune response (Porro et al., 2020). In the present study, the plasma level of IL-10 was significantly lower in the EAE group than in the CFA group, with the mean level of IL-10 in the RR-EAE group approaching 3.76 pg/mL on day 17 (p<0.05; median 3.24 pg/mL) (**Fig. 4A**), 6.35 pg/mL on day 30 (p<0.01; median 5.49) (**Fig. 4B**), and 17.18 pg/mL on day 45 (median 16.77 pg/mL) (**Fig. 4C**). The level of IL-10 was lowest on day 17 (clinical score ∼3-4), in the peak phase of disease, compared to other time points (**Fig. 4**). The mean concentration of IL-10 was 9.09 ± 4.11 pg/mL for RR-EAE, compared to 27.43 ± 7.95 pg/mL for CFA, with a detection range of 0.46 to 33.87 pg/mL for EAE and12.38 to 95.88 pg/mL for CFA (**Table 1**). Similarly, the same downward trend was found in chronic-EAE, with mean IL-10 concentrations of 7.82 ± 1.71 pg/mL in the EAE group and 34.06 ± 18.68 pg/mL in the CFA group, with detections ranging from 1.92 to 14.80 pg/mL in the EAE group and from 5.07 to 114.40 pg/mL in the CFA group (**Fig. 5; Table 1**). The maximum decrease of IL-10 concentration in EAE mice was observed on day 45 (mean level 9.92 pg/mL; *p* < 0.001; median 8.91 pg/mL) compared to day 20 (5.73 pg/mL; median 6.74 pg/mL). Overall, IL-10 concentrations decreased in both models of EAE.

**Fig 4.**
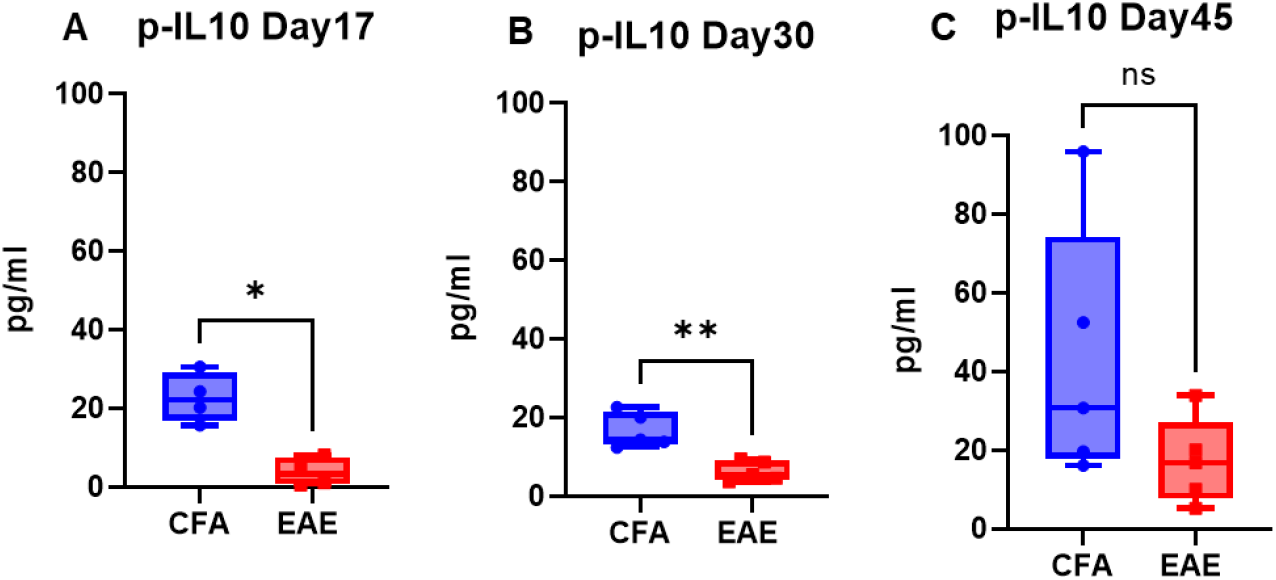
IL-10 profile in plasma of RR-EAE. Plasma level of IL-10 at respective time points in CFA vs EAE. Values are shown in pg/mL. *p<0.05, **p<0.01 (as determined by Mann-Whitney test).

**Fig 5.**
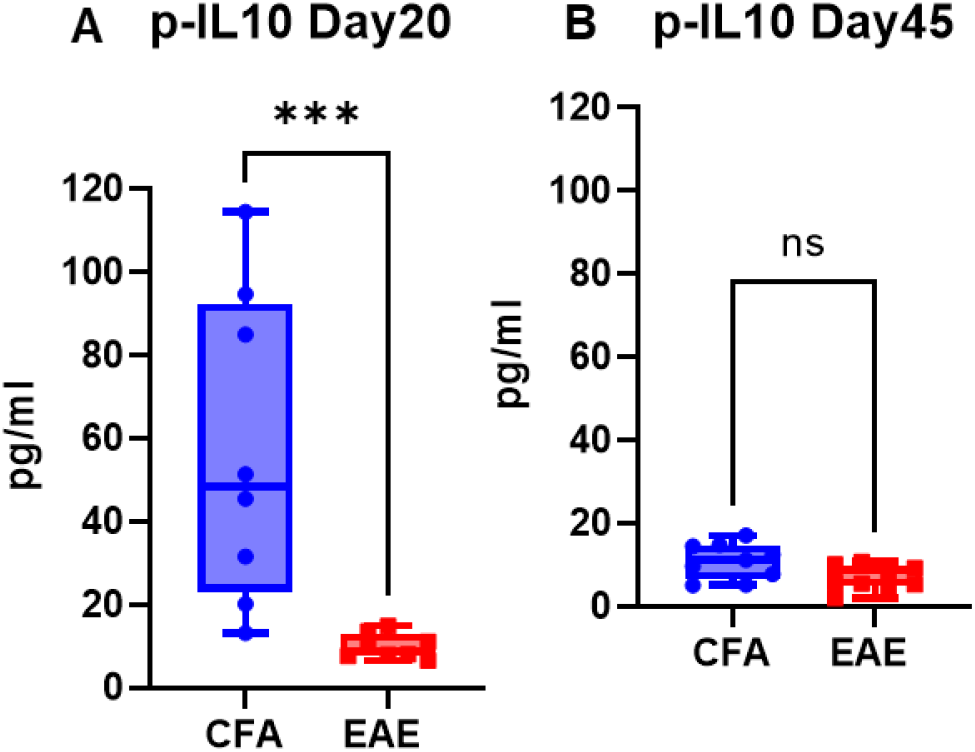
IL-10 profile in plasma of Chronic-EAE. Plasma level of IL-10 at respective time points in CFA vs EAE. Values are shown in pg/mL. ***p<0.001, ns non-significant (as determined by Mann-Whitney t-test) vs EAE.

**Fig 6.**
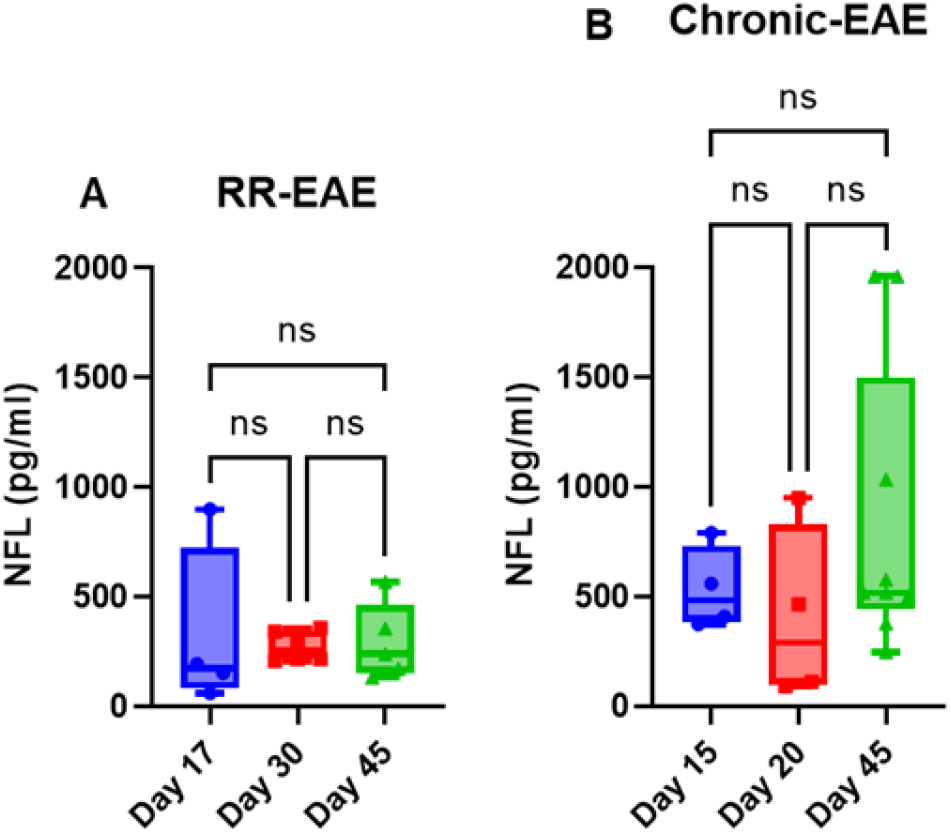
Comparative plasma NFL concentration along disease duration in RR-EAE and Chronic-EAE. Values are shown in pg/mL as determined by one-way ANOVA Kruskal-Wallis test, CFA vs EAE.

## 4. Discussion

The current goal of MS biomarker research is to use sensitive platforms that generate a specific biomarker profile to improve the diagnosis and management of MS. SIMOA technology is a fully automated platform that detects analytes (mainly proteins) when bound to antibody-coated beads combined with high-resolution fluorescence imaging (Rissin et al., 2010). Currently, the two most promising biomarker candidates for assessing neural damage in MS patients are NFL and GFAP; however, results are inconsistent across studies, which is highly suggestive of the variability in techniques used to measure them, as well as the heterogeneity across study cohorts (Arroyo et al., 2023). NFL represents the cytoskeletal component of neurons, while GFAP represents the cytoskeletal intermediate filament of astrocytes, and both serve as biomarkers of neuronal death, axonal degeneration, and astrogliosis (Yang & Wang, 2015). Blood NFL is a neuro-axonal injury marker associated with MS relapse, worsening of EDSS scores, lesions on MRI scans, and atrophy of the brain and spinal cord in patients with MS (Barro et al., 2018). NFL has been investigated as a potential prognostic and disease activity marker, with a potential relationship between its leakage and the rate of neurodegeneration (Bridel et al., 2019). Blood-based detection of NFL highlights early demyelination, while GFAP is mainly analyzed for MS progression and severity. The ability to measure NFL at different time points through serum detection also makes it suitable for monitoring treatment response. However, there are several limitations in the application of NFL detection. As a cytoskeletal protein that can be released by any kind of brain injury, it is not specific to MS; thus, any other neurologic disease or injury can have confounding effects. It is more logical to detect a panel of biomarkers using an ultrasensitive platform that would enhance the specificity, accuracy, and reproducibility of characterizing MS and post-treatment assessments.

The CNS biomarkers NFL and GFAP are used to monitor pathophysiological and neurodegenerative manifestations in preclinical MS and EAE animal models. The present study covers both RR-EAE (SJL) and chronic-EAE (B6) models, contrasting previously published reports that use only the B6 model (Gnanapavan et al., 2012; Aharoni et al., 2021). Since NFL leakage is associated with impaired blood-brain barrier integrity, immune cell extravasation, and CNS inflammation following the first demyelinating event in MS (Uher et al., 2021), we measured its plasma level in both EAE models. Notably, NFL leakage in the EAE model was more pronounced for the chronic-EAE model than the RR-EAE model, with no conclusive trend for kinetics across time points in either model. This contrasts the findings of Aharoni et al., who reported the greatest leakage to occur at the peak of disease followed by a subsequent decline as the disease progressed (Aharoni et al., 2021). However, this study was underpowered (n < 3) and variable across time points, limiting the significance of their data. Additionally, findings reported by Gnanapavan et al. showed opposite trends in B6 mice, possibly due to either the application of non-SIMOA methods for NFL detection, or batch variation in mice (Gnanapavan et al., 2012). In the present study, neurodegeneration in the form of neuroaxonal damage was confirmed by substantial leakage of NFL into peripheral circulation on day 17 (mean value 326.42 pg/mL) after disease induction in the RR-EAE model, overlapping with a maximum clinical score of ∼3-4 in these mice. As the disease progresses, NFL levels declined in EAE models at other time points (day 30 and day 45). Furthermore, the average level of NFL was greater in the chronic- EAE (B6) model (598.11 ± 133.83 pg/mL) compared to the RR-EAE (SJL) model (297.45 ± 15.77 pg/mL) for the combined time points, with the highest value in B6 nearing 1960.59 pg/mL. These findings further substantiate NFL as a reliable biomarker for assessing disease activity in EAE. However, the dynamics of NFL in the blood are a limiting factor, making it difficult to link NFL leakage with the degree of neurodegeneration and the extent of CNS damage. Similarly, we detected an increase in plasma GFAP after EAE induction in both models, with a maximum increase detected in the RR-EAE group (203.56 ± 173.60 pg/ml). Taken together, our findings for NFL and GFAP confirm the substantial leakage of CNS components into the periphery as a pathological manifestation of CNS damage in EAE.

Moreover, pathogenic cytokines (IFNγ, IL-17, TNFα, IL-6, IL-12p70, and IL-10) can also reflect the degree of inflammation in patients, as MS therapies are largely aimed at lowering the inflammatory response by modulating the levels of inflammatory markers (Wagner et al., 2020). Cytokine profiles have been established in various studies of adult and pediatric patients with MS (Bhise et al., 2016; Chen et al., 2012; Chen et al., 2012; Imitola et al., 2005; Martins et al., 2011); however, this process is far more complex since it involves an array of cytokines that have temporal effects on MS. Due to this complexity, detecting multiple variables, including CNS damage biomarkers, provides a more meaningful option for monitoring disease evolution. In the present study, the concentrations of cytokines IL-6, IL-17, IL-12p70, and TNFα were below the detection limit (conc. lower than the lowest calibrator) across all time points in most of the samples, which is why there was no conclusive data, making it difficult to interpret the trend in EAE (**Table 1**). However, we found a lower level of IL-10 in both models of EAE, with a mean of 9.09 ± 4.11 pg/mL in RR-EAE and 7.82 ± 1.71 pg/mL in chronic-EAE. Given the central role of IL- 10 in the pathophysiology of MS and other neurodegenerative diseases, its lower production in humans is considered a risk factor for MS (Porro et al., 2020; Vandenbark et al., 2001). IL-10 also regulates inflammation-mediated CNS damage in EAE, as evidenced by the exacerbation of EAE in IL-10 KO mice, compared to mice overexpressing IL-10 (Bettelli et al., 1998; Cua et al., 1999). Overall, our findings indicate that IL-10 is suppressed in EAE, confirming its anti-inflammatory role that suppress MS.

Notably, GFAP and IL-10 were not detected by SIMOA at the other time points mentioned in the study design, as the values for CFA and EAE mice in both models were either zero for GFAP or below the detection limit for IL-10. These readings could be due to machine error, the assay did not work for those samples, or issues with bead–analyte binding. Given the sensitivity of the SIMOA platform, sample processing was uniform across all batches, ruling out its impact on the data. Even so, our data indicated that disease severity parameters and plasma profiles of CNS biomarkers (NFL and GFAP) and IL-10 can be used together to monitor pathological manifestations over disease duration and to test therapeutic interventions in EAE models. These parameters can also be linked with other analyses to provide a comprehensive profile of drug testing in EAE. This finding was confirmed in our prior study showing a protective effect of the pro-resolution lipid mediator maresin1 (MaR1) in EAE, where we used SIMOA along with CNS-related histopathological data and other immunological parameters to validate this. Compared to those in the untreated group, plasma levels of NFL in MaR1-treated EAE-induced mice were lower, confirming the neuroprotective effect of MaR1 (Zahoor et al., 2025). Notably, we have also performed SIMOA assays in human patient samples, which again confirmed the use of NFL and GFAP as biomarkers of disease activity in MS, validating the utility of the SIMOA platform from animal models to human samples (Zahoor et al., 2022). The present study is the first to report multiple biomarker profiling in plasma throughout the disease-course of EAE using an ultrasensitive platform in two separate EAE models (Aharoni et al., 2021). One limitation of our study is it’s exclusivity to female mice, with a small sample size. However, we have used more than one EAE model (RR and Chronic) to show that any observed effects are pervasive.

## 5. Conclusions

Taken together, our results validate the utility of the SIMOA as a robust analytical platform with high sensitivity that can precisely detect small changes in samples, making it a highly valuable approach in clinical settings for both human patients with MS and EAE animal models.

At the same time, inconsistency in the data, variable sample size, and variation across mouse strains/batches due to unknown confounding variables cannot be ruled out, highlighting the use of caution when interpreting data across studies and making associations between these analytes and disease status. However, further large-scale prospective studies are warranted to validate the detection range and utility of these translational biomarkers in clinical practice.

## Funding

This work was supported in part by research grants from the National Multiple Sclerosis Society (US) (RG-1807-31964 and RG-2111–38733), the US National Institutes of Health (NS112727 and AI144004), and Henry Ford Health Internal Support (A10270 and A30967) to SG. The funders had no role in the study design, data collection, or interpretation or in the decision to submit the work for publication.

## Author Contributions

IZ designed the study, performed the assays, data analysis, and compiled the manuscript. SM reviewed and edited the manuscript. SG directed the study, reviewed the data analysis, and edited the manuscript before approval for submission.

## Informed Consent Statement

Informed consent was obtained from all subjects involved in the study.

## Data Availability Statement

All the data generated or analyzed during this study are included in this published article.

## Conflicts of Interest

The authors declare that they have no conflicts of interest.

## Ethics Approval

The animal studies performed in this manuscript were approved by the IACUC committee of Henry Ford Health. The article does not contain any studies with human participants.

## Acknowledgements

The authors thank Katie H. Hurrish, Ph.D. at Henry Ford Health for language and editorial assistance.

## Notes

### Competing Interest Statement

The authors have declared no competing interest.

### Summary of Updates

The revised version has more detailed information.

